# Optimized AAV to express the unfolded protein response transcription factor XBP1s ameliorates Alzheimer’s disease features in mouse models

**DOI:** 10.64898/2026.08.11.743979

**Authors:** Mei Li Diaz, Giovanni Tamburini, Diego Arriagada, Natalia Poblete, Álvaro O. Ardiles, David Neira, Denisse Sepulveda, Gabriela Martínez, Romina Gozalvo, Javiera Arcos, Claudia Sepulveda-Quiñenao, Els Henckaerts, Sergio T. Ferreira, Adrian G. Palacios, Claudio Hetz

## Abstract

Proteostasis impairment at the level of the endoplasmic reticulum (ER) is a salient feature of Alzheimer’s disease (AD). The unfolded protein response (UPR) is the main pathway to cope with ER stress, where the expression of the transcription factor X-Box binding protein 1 (XBP1) is central to establish repair programs. To artificially enforce the adaptive capacity of the UPR in the AD brain, we recently reported the protective effects of overexpressing active XBP1 in the brain using adeno-associated vectors (AAVs) of AD mice, in addition to aged animals. Here we have generated a next generation vector suitable for clinical testing by (i) expressing codon-optimized human XBP1s without artificial tags, (ii) the use of the synapsin promoter to restrict expression to neurons, and (iii) incorporating a novel variant of AAV2 (AAV-TT) with greater biodistribution (here termed Proteostaser-1). Treatment of 5xFAD mice with Proteostaser-1 improved spatial learning and synaptic plasticity, and reduced the deposition of amyloid plaques in the brain. Proteostaser-1 administration also improved cognition in a model of sporadic AD based on the intracerebral injection of amyloid β oligomers. Our results further support the therapeutic potential of the UPR as a strategy to ameliorate AD features and sustain synaptic function.

## INTRODUCTION

Alzheimer’s disease (AD) is the most common form of dementia in the elderly, involving progressive synaptic dysfunction, neurodegeneration, and cognitive impairment ^1,2^. AD is characterized by the abnormal deposition of protein aggregates in the brain formed by amyloid β and hyperphosphorylated Tau ^3^. Mutations in the genes encoding APOE, presenilin 1, presenilin 2, and amyloid β precursor protein (APP) are causative factors of AD ^4^. Nevertheless, aging is still the main risk factor for developing AD, with the majority of patients presenting clinical symptoms after the age of 65. However, the functional relationship between the biology of aging and the risk of developing AD remains poorly understood.

The decline in the buffering capacity of the proteostasis network has been identified as a primary hallmark of aging ^5,6^, a phenomenon that may contribute to AD pathogenesis. Proteostasis is maintained through the dynamic integration of the pathways that mediate the synthesis, quality control, folding, degradation, and targeting of proteins to their destination^7^. One of the central nodes of the proteostasis network altered during aging involves the function of the endoplasmic reticulum (ER), the main site of protein production in the cell ^8^. The ER is also highly altered in AD^9–11^. To cope with ER stress, cells activate an evolutionarily conserved pathway known as the unfolded protein response (UPR), which aims to re-establish proteostasis^12^. The UPR reinforces many processes involved in the function of the secretory pathway to improve protein production and sustain cell function, whereas chronic ER stress results in neurodegeneration and cell death^13^. The most conserved UPR signaling branch is initiated by the ER stress sensor inositol-requiring enzyme 1α (here referred to as IRE1). IRE1 catalyzes the unconventional splicing of the mRNA encoding X-box binding protein-1 (XBP1), excising a 26-nucleotide intron^14^. This processing event shifts the XBP1 mRNA open reading frame, resulting in the expression of an active transcription factor, termed XBP1s, that operates as a master regulator of UPR transcriptional reprogramming. We recently reported that the activity of the IRE1/XBP1 pathway dictates normal brain health span^15^. Strategies to express XBP1s in neurons, using either transgenic mice or gene therapy, delayed synaptic dysfunction and cognitive decline during normal aging, in addition to reducing the content of senescent cells in the brain^15^. XBP1s expression in the hippocampus largely corrected the proteomic changes associated with aging, affecting the expression of a cluster of proteins involved in synaptic function, plasticity, and pathways linked to neurodegenerative diseases^15^.

Several studies have addressed the possible contribution of the UPR to AD. Overexpression of XBP1 in fly models of AD provided neuroprotection against amyloid β and Tau^16,17^. It was also reported that the behavioral impairment triggered by amyloid β in *C. elegans* is prevented by the overexpression of XBP1s, whereas knocking down *Xbp1* in worms exacerbates amyloid β pathogenesis^18,19^. Other studies reported that XBP1s expression increases amyloid β and Tau clearance^18–20^, suggesting multiple mechanisms of neuroprotection^18,21^. The endogenous activity of IRE1 is transiently induced in mouse models of AD^22^ and declines in AD fly models with age^16^, consistent with data reported in human post-mortem AD brain tissue^22^. Based on these findings, and because XBP1s is a transcription factor that regulates multiple nodes of the proteostasis network, we proposed that strategies to artificially enforce XBP1s-dependent responses may improve neuronal function in AD^23^. To test this concept, we recently demonstrated that the local delivery of mouse XBP1s into the hippocampus of AD mice using adeno-associated virus (AAV) improved long-term potentiation, memory performance, and dendritic spine density, in addition to reducing amyloid deposition^24^. Moreover, quantitative proteomics of the hippocampus revealed that XBP1s overexpression corrects a large proportion of the alterations in protein expression observed in AD mice, restoring the levels of several synaptic proteins and factors involved in neuronal morphology and axonal growth^24^.

Gene therapy is an active area of development to generate treatments for neurodegenerative diseases, based on the recent Food and Drug Administration (FDA) approval of AAV-mediated therapies for spinocerebellar ataxia, Duchenne muscular dystrophy, Leber’s congenital amaurosis, among others ^25^. Many clinical trials are under development in AD using AAV, already demonstrating favorable safety properties and possible improvements^26^. One of the main challenges in developing gene therapies for CNS diseases is the inefficient biodistribution of AAVs in the brain, as revealed by low viral spreading from the site of injection in postmortem studies from patients enrolled in a Phase II clinical trial in AD (AAV serotype 2 expressing NGF)^27^. To overcome these problems, a variety of capsid engineering methods have been used to generate recombinant viruses with improved spread and transduction properties at lower titers^28^. Using this approach, we previously generated an AAV capsid incorporating amino acids conserved among natural AAV2 isolates and tested its biodistribution properties in mice and rats^29^. Intriguingly, this novel variant, termed AAV-TT, demonstrated strong neurotropism in rodents and displayed significantly improved biodistribution throughout the central nervous system compared to regular AAV2^29^. AAV-TT may have lower antigenic activity and use non-glucan receptors for cell internalization^30^. In this study, we optimized our previous AAV2-mXBP1s/EGFP gene therapy ^24^ to generate an improved vector with features intended to facilitate future clinical development. To this end, we eliminated the EGFP cassette, used codon-optimized human *XBP1s* cDNA instead of the mouse gene, replaced the constitutive promoter with a synapsin-1 mini promoter, and used an AAV-TT serotype for packaging. For simplicity, we termed AAV-TT-Syn-hXBP1s “Proteostaser-1”. Delivery of Proteostaser-1 into the hippocampus of 5xFAD mice improved cognition and synaptic function, as well as decreased amyloid β load. In addition, Proteostaser-1 was protective in a model of sporadic AD based on the delivery of small amyloid β oligomers (AβOs) into the hippocampus. Our results reinforce the concept that strategies to enhance the activity of adaptive UPR may be protective in AD.

## RESULTS

### Generation of an optimized viral construct to deliver XBP1s into the central nervous system

The human *XBP1s* cDNA sequence without any tag or reporter cassette was designed with codon-optimized sequences. This cDNA was synthesized and inserted into an AAV plasmid under the control of the synapsin-1 mini promoter to drive expression in neurons (Figure S1A). This construct was then packaged into AAV-TT, a modified AAV2 serotype, and purified and quantified (see Materials and Methods). We then carried out experiments to confirm expression in the hippocampus of mice and compared it to the previously characterized AAV2-mXBP1s/EGFP (mouse cDNA) construct, which uses the CMV promoter for transgene expression and also contains an ELF2 EGFP cassette for detection ^31^. Two-month-old animals were bilaterally injected into the hippocampus using stereotaxis surgery, and one month later, the levels of human and mouse XBP1s mRNA were measured by quantitative PCR. Higher levels of expression were observed when Proteostaser-1 was injected compared to AAV2-mXBP1s/EGFP (Figure S1B). We obtained similar results when the expression of XBP1s was measured in total protein extracts from dissected hippocampus (Figure S1C). These experiments demonstrated that the use of Proteostaser-1 results in more than 3-fold higher expression than AAV2-mXBP1s/EGFP. Finally, we re-quantified the AAV viral genomes of AAV2-mXBP1s and AAV-TT-Syn-hXBP1s side by side, and the results indicated that the viral particles used for AAV2-mXBP1s/EGFP had 4.5-fold higher concentrations than Proteostaser-1 (Figure S1C). Taken together, these results demonstrate a greater expression efficacy of XBP1s in the brain when Proteostaser-1 particles are injected compared to AAV2-mXBP1s/EGFP.

### Expression of XBP1s in the hippocampus reduces amyloid β load in AD mice

To evaluate the therapeutic potential of Proteostaser-1 in preclinical models of AD, we locally targeted the hippocampus using bilateral stereotaxic injections. 5xFAD mice were injected with Proteostaser-1 or an equal volume of PBS as control (Sham) at 8 weeks of age, and their behavioral performance was then monitored at different ages. This AD mouse model expresses a combination of five human mutations in the APP and presenilin-1 (PSEN1) genes and accumulates amyloid plaques from approximately 2 to 4 months of age, resulting in learning and memory deficits by 6 months of age ^32^. Immunohistochemical analysis of amyloid β content (total number of plaques per area) using the 4G8 antibody in animals at 6 and 9 months of age indicated a significant reduction in amyloid β deposition in the hippocampus of Proteostaser-1-treated 5xFAD mice compared to sham 5xFAD controls at both time points (Figure 1A and 1B, respectively). Amyloid β burden (percentage of area occupied) was significantly reduced at 9 months (Figure 1B), whereas at 6 months plaque number, but not burden, differed between groups (Figure 1A). Next, we assessed fibrillar amyloid deposits using thioflavin S (ThS) staining in hippocampal CA1 sections from 9-month-old 5xFAD mice. Consistent with our immunohistochemical analysis, ThS staining confirmed a significant reduction in fibrillar amyloid deposition in the hippocampus of Proteostaser-1-treated mice compared with PBS-treated controls (Figure 1C). In addition, we extended our analysis to the cerebral cortex, another brain region severely affected during AD progression, and observed a similar reduction in ThS-positive amyloid deposits following Proteostaser-1 administration (Figure 1D). Together, these findings demonstrate that hippocampal overexpression of human XBP1s reduces amyloid β load in 5xFAD mice.

**Figure 1.**
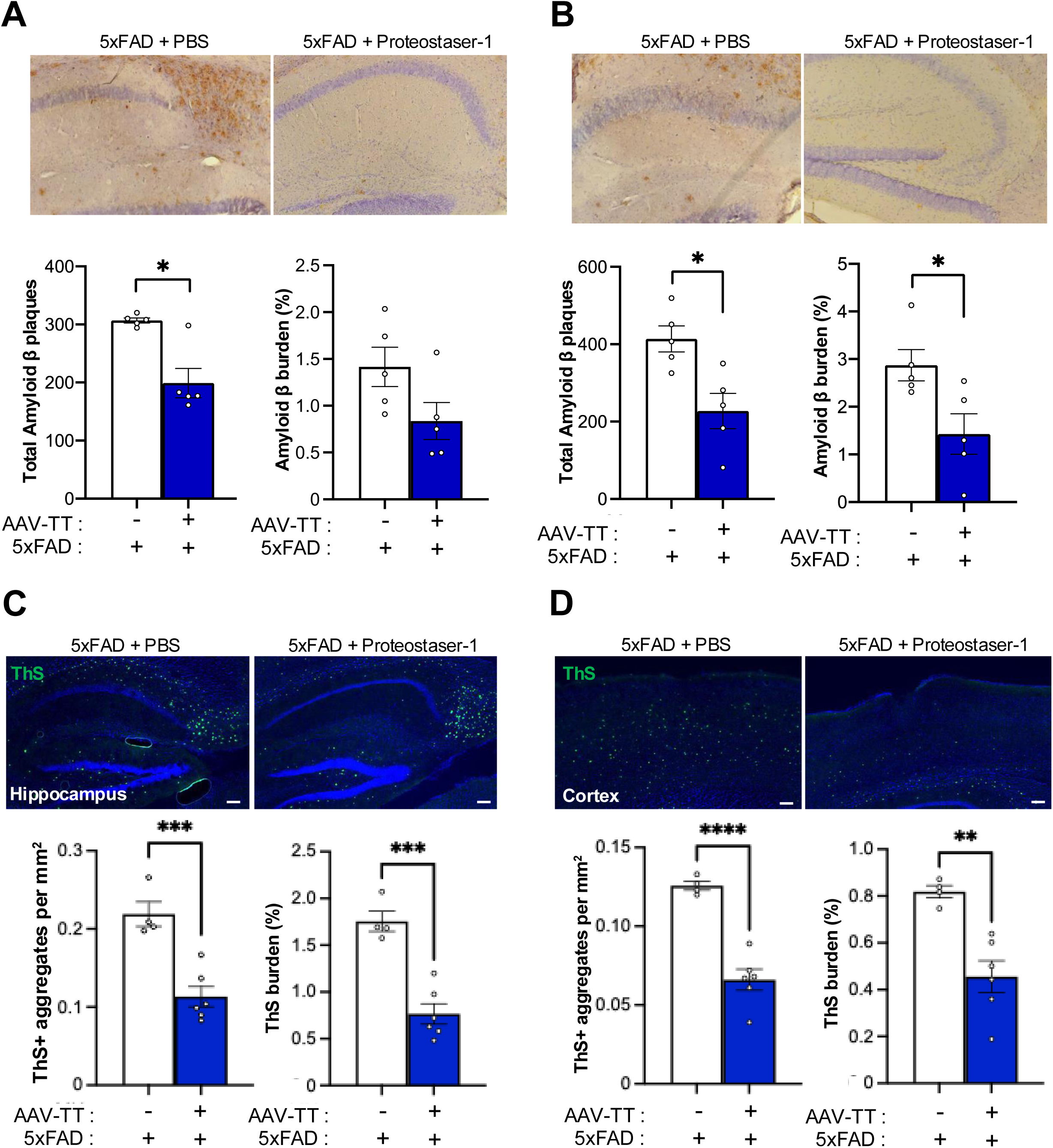
Delivery of Proteostaser-1 into the hippocampus of 5xFAD mice reduces amyloid β deposition. 5xFAD mice were injected with PBS or Proteostaser-1 (AAV) by bilateral stereotaxic injection at 2-months of age. Representative images of hippocampal CA1 sections are shown for immunohistochemical detection of amyloid β plaque deposition in 5xFAD mice treated with PBS or Proteostaser-1 at 6 months (A) and 9 months (B) of age. The total number and the percentage of area occupied by amyloid plaques were quantified (n = 5 per group). Representative images of thioflavin S (ThS) staining to monitor fibrillar amyloid β deposits in brain sections from 9-month-old 5xFAD mice treated with PBS or Proteostaser-1. ThS-positive signals were evaluated in the hippocampus (C) and cerebral cortex (D), and quantification for ThS-positive aggregates per area (mm^2^) and ThS-positive percentage was performed (n = 4 control; n = 6 AAV-TT group; bottom pannels). Mean and standard error are shown. Statistical analysis was performed using the Mann-Whitney U test. *: *p* < 0.05; **: *p* < 0.01; ***: *p* < 0.001; ****: *p* < 0.0001; ns, not-significant. Scale bar = 100 μm.

### Administration of Proteostaser-1 into the hippocampus of 5xFAD mice improves cognitive performance

Next, we determined the possible beneficial effects of Proteostaser-1 on the learning and memory capacity of 5xFAD animals. A cohort of mice received bilateral injections of Proteostaser-1 or PBS into the hippocampus at 2 months of age, followed by analysis in 6-months-old animals. Analysis of the learning curve in the Morris water maze indicated a significant decrease in the latency to find the hidden platform in Proteostaser-1-treated 5xFAD mice compared to sham controls (Figure 2A). In addition, assessment of thigmotaxis, measured as the distance traveled in the absence of the hidden platform, indicated lower distance traveled in 5xFAD mice when Proteostaser-1 was administered (Figure 2B, left panel, and Figure S4), suggesting a decrease in anxious behavior. Analysis of long-term thigmotaxis did not show significant differences between treatments in 5xFAD animals (Figure 2B, right panel). We also performed a second behavioral test, the Barnes maze, in animals treated with Proteostaser-1 using an independent cohort. We observed an improvement in the learning curve of 9-months-old animals treated with Proteostaser-1 compared to sham controls (Figure 2C), whereas short-and long-term memory retention measured on the test days did not differ between treated and sham 5xFAD animals (Figure 2D). Thus, local delivery of AAV-TT expressing XBP1s into the hippocampus partially ameliorates the learning and anxiety deficits observed in our AD model.

**Figure 2.**
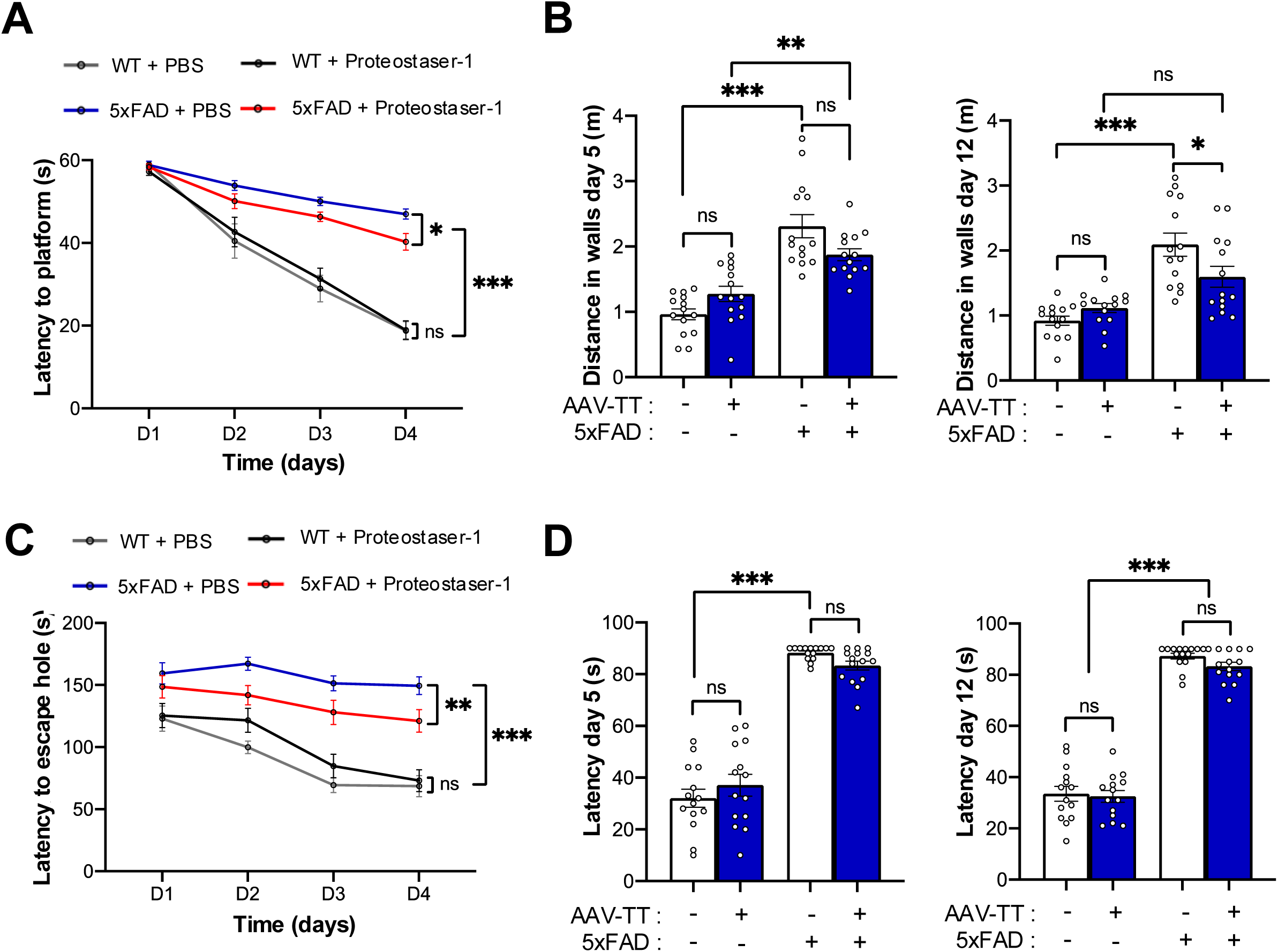
Proteostaser-1 treatment ameliorates cognitive deficits in 5xFAD mice. 5xFAD mice or littermate non-transgenic animals were injected in the hippocampus with PBS or Proteostaser-1 by bilateral stereotaxis injection at 2-months of age. (A) At 6 months of age, memory performance was assessed using the Morris water maze. Escape latency to reach the platform was measured in WT mice injected with PBS (n = 15), WT mice injected with Proteostaser-1 (n = 15), 5xFAD mice injected with PBS (n = 15), and 5xFAD mice treated with Proteostaser-1 (n = 15). (B) Thigmotaxis analysis was performed on the testing days (days 5 and 12). (C) A second cohort of animals was evaluated using the Barnes maze as indicated in A. Learning performance was assessed as latency to reach the target hole. (D) Short- and long-term memory were evaluated based on latency to reach the target hole on the test day. Experimental groups included WT mice injected with PBS (n = 14), WT mice injected with Proteostaser-1 (n = 15), 5xFAD mice injected with PBS (n = 15), and 5xFAD mice treated with Proteostaser-1 (n = 15). Mean and standard error are shown. Statistical analysis was performed using two-way ANOVA followed by Tukey’s post hoc test. *: *p* < 0.05; **: *p* < 0.01; ***: *p* < 0.001; ns, not-significant.

### Proteostaser-1 administration into the hippocampus of AD mice restores synaptic plasticity

Next, we measured long-term potentiation (LTP) *ex vivo* in hippocampal slices obtained from animals of 6 or 9 months of age that were treated or not with Proteostaser-1. Analysis of the youngest group showed considerable variability and no significant differences between groups (Figure 3A); with five animals per group this experiment is powered only for large effects and is therefore inconclusive rather than negative. In contrast, when 9-month-old animals were analyzed, a significant improvement in LTP was observed when Proteostaser-1 was delivered into the hippocampus of 5xFAD mice compared to PBS-treated 5xFAD mice (Figure 3B).

**Figure 3.**
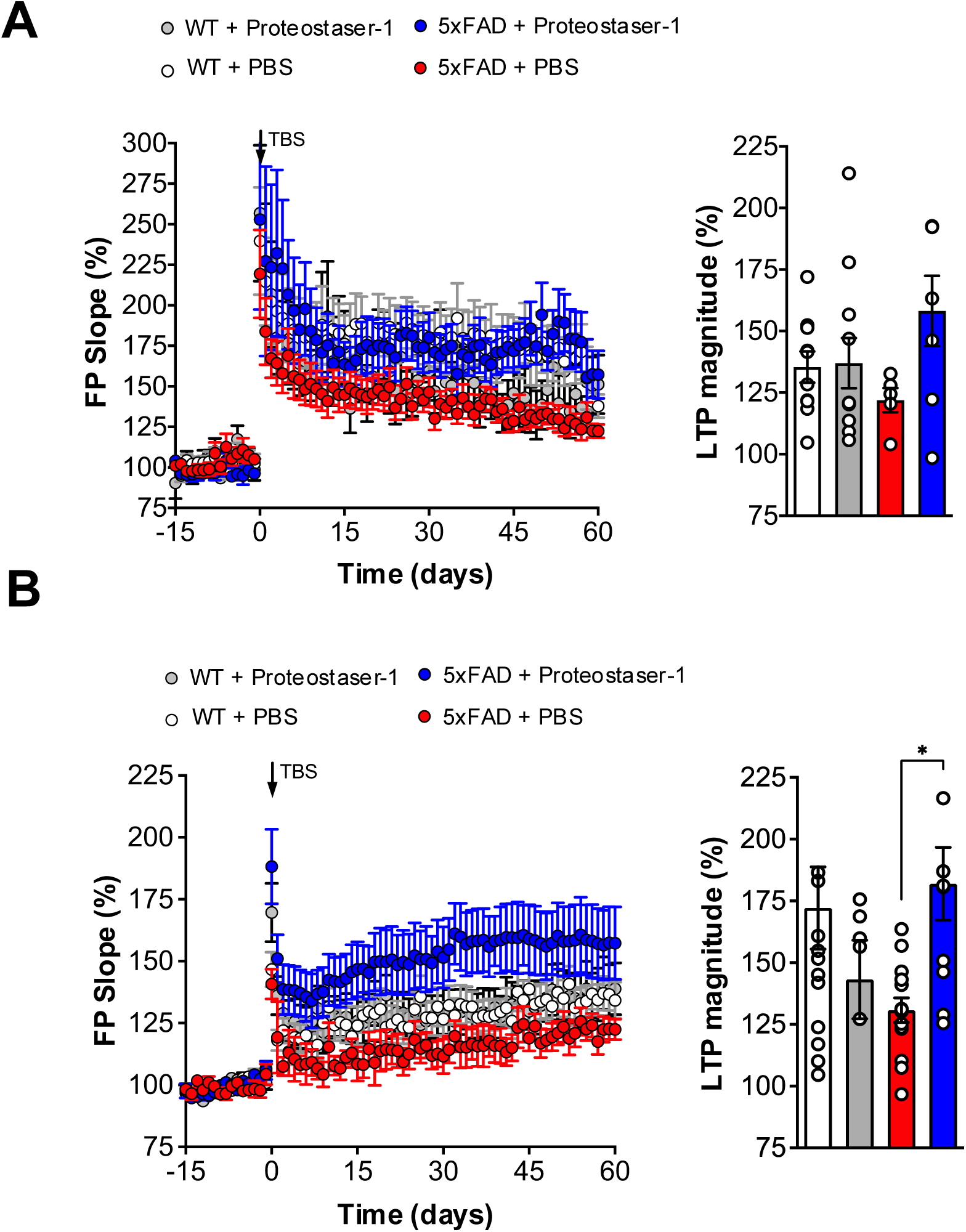
Proteostaser-1 treatment restores hippocampal long-term potentiation in 5xFAD mice. 2-month-old 5xFAD mice or non-transgenic littermate control animals were injected with PBS or Proteostaser-1 by bilateral stereotaxis injection into the hippocampal region. Long-term potentiation (LTP) was then measured *ex vivo* in hippocampal slices isolated from animals of 6- (A) or 9-months of age (B). LTP was induced using a high-frequency theta-burst stimulation protocol, applied after a stable 15-minute period of synaptic transmission and recorded for up to 60 minutes after stimulation. Right panel shows the average LTP magnitude during the final minutes of recording (n = 5 per group). Mean and standard error are shown. Statistical analysis was performed using the Mann-Whitney U test. *: *p* < 0.05; **: *p* < 0.01; ***: *p* < 0.001; ns, not-significant.

### Proteostaser-1 provides protection against AβO administration

To validate our technology in a model that does not rely on overexpression of mutant transgenes and instead resembles molecular aspects occurring in sporadic cases, we injected non-toxic doses of synthetic AβOs into the hippocampus, which generate a rapid but transient impairment in synaptic function ^33,34^, followed by cognitive assessment. Mice were injected with Proteostaser-1 or PBS, and 30 days later underwent a second surgery to infuse 20 pmol of AβOs (Figure 4A). After a 5-day recovery period, cognitive assessment was performed in the Morris water maze. Animals injected with the AβOs showed robust and significant memory impairment, reflected in increased latency to find the platform, a phenotype completely reversed by administration of Proteostaser-1 (Figure 4B). In addition, animals injected with AβO showed a significant decrease in the distance traveled in the quadrant occupied by the platform during both the short- and long-term memory tests; Proteostaser-1 prevented this deficit in the short-term test only, and the long-term deficit persisted in treated animals (Figure 4C). Overall, this alternative model allowed us to confirm the beneficial effects of Proteostaser-1 on spatial learning in AD models.

**Figure 4.**
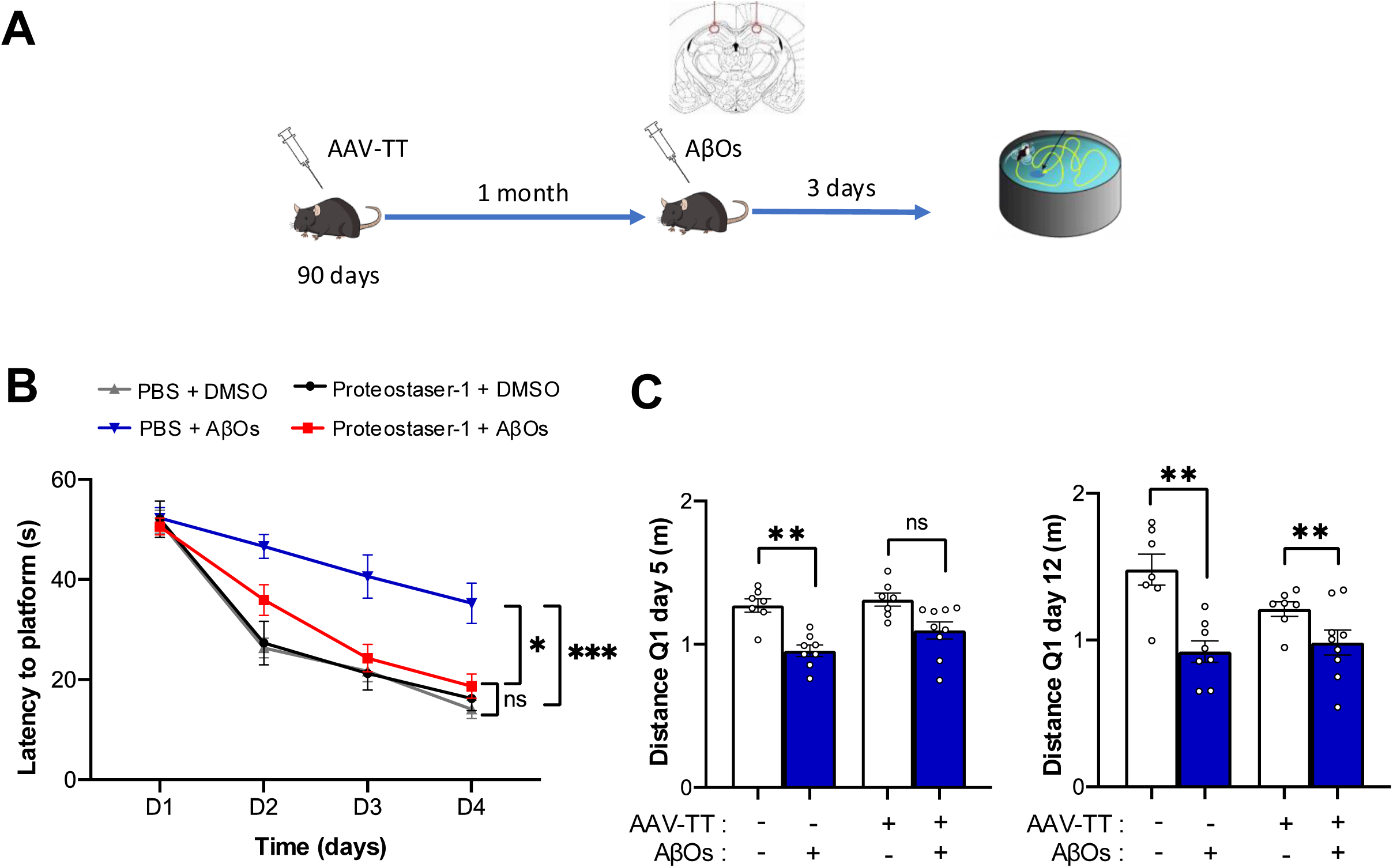
Hippocampal delivery of Proteostaser-1 improves cognitive performance in animals injected with AβOs. WT mice of 90 days of age were injected with PBS or Proteostaser-1 into the hippocampus by bilateral stereotaxic injection. One month later, mice received an injection of 20 pmol of synthetic AβOs (see material and methods) at the same injection site, and behavioral evaluation was performed 3 days after AβOs administration. (A) Experimental timeline showing bilateral hippocampal injection of PBS or Proteostaser-1, followed by AβOs administration and subsequent behavioral assessment. (B) Cognitive performance was assessed using the Morris water maze. Learning performance was evaluated as escape latency to reach the platform (n = 7–9 per group). (C) Short- and long-term memory were evaluated based on the distance traveled in the target quadrant (n = 7–9 per group). Mean and standard error are shown. Statistical analysis was performed using the Mann-Whitney U test. *: *p* < 0.05; **: *p* < 0.01; ***: *p* < 0.001; ns, not-significant.

## DISCUSSION

According to The World Alzheimer Report 2021 (Gauthier S.; R.-N.P., 2021), approximately 55 million people live with dementia worldwide. Current FDA-approved treatments provide, at best, mild symptomatic relief, and with no disease-modifying therapy in sight, AD is perhaps the most poignant example of a neurodegenerative disease with an unmet need. Thus, gene therapy is particularly attractive, since it promises to target disease etiology and provide long-term correction^35^. Attenuation of ER stress with pharmacological or gene therapy strategies affords strong protection in various animal models of brain diseases, and thus the pathway holds promise as a target for developing therapeutics for human neurodegenerative diseases, including AD^36–38^. AAVs are the most common vectors for gene therapy of the CNS, mainly due to their safety and efficacious neuronal transduction^25^. Nevertheless, clinical efficacy for AD indications remains limited ^26,39^, and one reason for this is the relatively limited spread and transduction efficacy in large regions of the human brain.

Here we optimized a research tool to artificially enforce UPR-dependent repair programs into an AAV prototype that could be tested in the clinic in the near future. Our results indicate that Proteostaser-1 results in better expression of XBP1s compared with the previously described AAV2 construct. Proteostaser-1 administration ameliorated AD-related phenotypes in independent animal models. The four modifications were introduced together, so the present data cannot attribute the improvement to any single one; we propose that the use of the AAV-TT capsid is an important optimization, because AAV2 has been reported to have lower efficiency as a vector for the CNS compared with other AAV serotypes, attributed to its broad tissue tropism due to a high affinity for heparan sulfate proteoglycan (HSPG)^40^, whereas AAV-TT has a reduced HSPG-binding site ^29,30^. However, studies in monkeys compared AAV-TT with AAV9 (not AAV2) and suggested only slight enhancement of viral spreading through the brain^41^. Another factor contributing to gene therapy effectiveness and safety is the promoter, which influences the strength and cell selectivity of transgene expression^42^. In this regard, a study aiming to identify promoters for optimal transduction in the retina found that eGFP expression was strongest in the retina, optic nerves, and brain when the human synapsin promoter was used compared to cytomegalovirus^43^, supporting the selection of this promoter in the current study.

Accumulating evidence supports the concept that impairment in the adaptive and buffering capacity of the proteostasis network is a salient feature of AD, in addition to aging, the major risk factor for developing AD^44^. The involvement of protein folding stress responses at the ER is a transversal pathological event observed in AD patient-derived tissue, as well as in most animal and cellular models of the disease^9,45,46^. Importantly, XBP1s is a key transcription factor of the UPR that mediates adaptive responses through the upregulation of multiple genes involved in proteostasis mechanisms, in addition to genes involved in intrinsic neuronal functions. We reported that the beneficial effects of XBP1s expression in the context of experimental AD and normal aging involve a substantial correction of gene expression patterns associated with synaptic function, neuronal morphology, and connectivity^15,24^. More recently, we demonstrated that AAV-XBP1s administration also protects models of frontotemporal dementia, associated with cognitive improvement^47^. In this line, recent reports suggested that differential XBP1s expression in fly models of neurodegeneration may underlie the selective neuronal vulnerability observed^48^. In agreement with this concept, we report here that delivery of Proteostaser-1 into the hippocampus of AD mice reduces amyloid content and improves synaptic function and cognitive performance, suggesting a possible impact at the level of proteostasis and neuronal physiology. On this line, XBP1s has alternative roles in the nervous system, regulating BDNF expression and memory function^49,50^, and our studies in models of normal brain aging and AD using quantitative proteomics indicated that the major effects of XBP1 in gene expression are associated with synaptic function, neuronal morphology and connectivity^24,51^. We therefore speculate that a major protective mechanism of XBP1s in AD relates to its function as a regulator of neuronal physiology, although further research may elucidate the underlying mechanisms of action. In agreement with our results, Cissé et al. reported that brain delivery of XBP1s using lentiviruses improved synaptic plasticity and cognitive function in 3xTg-AD mice, a less aggressive AD model compared to 5xFAD, through regulation of the synaptic regulator Kalirin-7 ^50^.

The artificial enforcement of XBP1s-dependent responses using gene therapy has been reported to provide beneficial effects in many different disease models. For example, ectopic expression of XBP1s improved dopaminergic neuron survival in Parkinson’s disease ^52,53^, reduced mutant huntingtin aggregation in the striatum^54^, improved locomotor recovery after spinal cord injury^31^, accelerated axonal regeneration of peripheral nerves^55^, and enhanced synaptic plasticity at basal levels^49^. Hundreds of clinical trials have already demonstrated the safety and applicability of AAV-mediated gene therapy for treating human diseases. By early 2025, there were already more than 340 active clinical trials involving AAV vectors. Currently, there are at least seven AAV-based gene therapies approved by the FDA/EMA for human use, including four technologies targeting nervous system disorders: Luxturna® for Leber congenital amaurosis, Zolgensma® for spinal muscular atrophy, Elevidys® for Duchenne muscular dystrophy, and Kebilidi®, which treats a rare neurodevelopmental disorder. In recent years, significant effort has been directed toward developing gene therapies to modify the course of AD using intracerebral injection of AAVs or intrathecal infusion of ASOs. Additionally, *ex vivo* gene therapies for AD have been employed to modify autologous fibroblasts for the delivery of trophic factors to the brain^56,57^. Key targets investigated in clinical trials for AD include nerve growth factor (NGF), brain-derived neurotrophic factor (BDNF), apolipoprotein E2 (APOE2), and human telomerase reverse transcriptase (hTERT)^26^. All these therapies currently in clinical phases are based on robust data derived from preclinical animal models. Finally, we would like to highlight that the use of transcription factors for gene therapy received FDA clearance (IND) to begin clinical testing January 2026, for trial conducted by Life Bioscience. This clinical trial (NCT07290244) aims to mitigate the adverse effects of aging on eye function. The trials will employ the ER-100 gene therapy, which involves the regulated expression of three transcription factors (OCT-4, SOX-2, and KLF-4-the Yamanaka factors) via AAV vectors, controlled through drug administration. These developments reinforce the viability of the concept behind Proteostaser-1, wherein transcriptional reprogramming represents a robust, cutting-edge therapeutic avenue that targets gene regulatory networks rather than specific genes or enzymes, theoretically yielding more robust and functional cellular effects. There remain multiple challenges in improving current gene therapies undergoing clinical and preclinical development^58^. Various challenges and limitations persist, such as optimizing gene delivery methods, improving safety and efficacy profiles, managing large-scale production costs, and determining long-term outcomes. Current strategies include engineering AAV capsids to facilitate blood-brain barrier penetration, reduce peripheral immunogenicity, and enable the delivery of multiple transgenes to address the complex pathology of AD and other neurodegenerative diseases. In summary, gene therapy has shown outstanding efficacy in treating various diseases in clinical trials; it holds the possibility of intervening pathogenic pathways or delivering disease-modifying agents, such as XBP1s, into the brains of AD patients. Preclinical studies in non-human primates are still needed to assess safety issues related to the chronic expression of adaptive UPR mediators. However, inducible and controlled systems could be further tested as designed in the ER-100 trials.

## MATERIALS & METHODS

### Design, Production, and Purification of AAV-XBP1s Vectors

The generation of AAV-XBP1s and AAV-Mock was described previously^29^. In brief, the entire murine *Xbp1s* expression cassette was excised from pcDNA3-XBP-1s as a MfeI/SphI fragment and inserted into a proviral plasmid, pAAVsp70, containing AAV inverted terminal repeats (ITRs). The *Xbp1s* cDNA is controlled by the CMV promoter. The vector also carries an EGFP expression cassette controlled by the ELF2 promoter, which serves as a fluorescent marker to report transduced cells. This vector is referred to as AAV-XBP1s, whereas the control vector contains a stuffer DNA sequence in addition to the EGFP cassette (AAV-Mock). Recombinant AAV-XBP1s (serotype 2/2) was produced by triple transfection of HEK293T cells using a rep/cap plasmid and pHelper (**Agilent Technologies, Santa Clara, CA, USA**), and purified by column affinity chromatography, as previously described ^59^. To obtain pure and concentrated AAV particles, cell lysates of transfected HEK293T cells were treated with trypsin and nuclease, followed by ion-exchange chromatography using ceramic hydroxyapatite and DEAE-Sepharose in combination with cellufine sulfate affinity chromatography. Viral titers were determined by real-time TaqMan PCR assay using specific primers for the bovine growth hormone (BGH) polyadenylation sequence, which is present in both plasmids.

### Animals

Wild-type C57BL/6 and 5xFAD animals (formerly JAX Stock No. 008730) were used. 5xFAD mice overexpress two transgenes, carrying three mutations in the human APP gene (Swedish, Florida, and London Familial Alzheimer’s Disease) and two mutations in the human presenilin-1 (PSEN1) gene (M146L+L286V). These animals begin to accumulate amyloid β aggregates at 2 months of age. Increased senile plaques, synaptic degeneration, gliosis, and cognitive impairment are observed between 4 and 5 months of age^60^. Animals were housed in groups of up to five in individually ventilated cages under standard conditions (22°C, 12 h light-dark cycle), receiving food and water *ad libitum*. Both sexes used and were balanced across experimental groups. Animals were allocated and image quantification were blinded to group allocation. Sample size was based on power analysis and previous studie^61^; no animals were excluded in behavioral studies. For histological analysis a subgroup of animals was assessed as indicated in figure legends. All animal manipulations were carried out according to standard regulations and were approved by the Animal Welfare Committee of the Faculty of Medicine of the Universidad de Chile, Chile (CBA 1141). Mice were euthanized by CO_2_ inhalation, and all efforts were made to minimize suffering. The right hemispheres were frozen at -80°C for biochemical analyses, whereas the left hemispheres were stored for histological studies.

### Behavioral studies

Cognitive impairment was measured using the Barnes maze and the Morris water maze. Briefly, the Barnes maze test consists of a circular surface containing 20 holes, one of which is leads to a hiding chamber (the target hole). Mice were placed on the maze surface and allowed to explore it for 3 minutes. Mice were exposed to an aversive sound stimulus to motivate escape into the hiding chamber. The room was equipped with spatial cues for mouse orientation. This training procedure was performed four times a day for four consecutive days per animal. On the fifth day, short-term memory was evaluated by removing the chamber; the animal was allowed to explore for 1.5 min, and latency to find the target hole was recorded. This trial was repeated on day 12 to assess long-term memory.

The Morris water maze was performed as previously described ^62^. In this assessment, animals learn to swim to a submerged, hidden platform. Mice at 6 or 9 months of age were placed in the pool and allowed to explore it for 1 min. The testing room was equipped with spatial cues for orientation. This training procedure was performed six times daily for four consecutive days per animal. The time spent by each animal to find the platform was measured during the training phase to evaluate learning performance. On day five, the platform was removed from the pool, and the time spent in the target quadrant was recorded to assess short-term spatial memory. Short- and long-term memory were measured on days 5 and 12, respectively; at the end of the test, a trial with a visible platform was conducted to exclude any visual or motor deficiency.

### Electrophysiological analysis

Hippocampal slices were prepared as previously reported ^49,61^. Mice at 6 to 9 months were deeply anesthetized with isoflurane, and their brains were quickly removed. Slices (350 μm) were dissected in ice-cold dissection buffer using a vibratome (Vibratome 1000 Plus, Ted Pella Inc., CA, USA). Synaptic responses were evoked by stimulating the Schaffer collaterals with 0.2-ms pulses delivered through concentric bipolar stimulating electrodes and recorded extracellularly in the stratum radiatum of the CA1 subfield, as described previously ^49^. Long-term potentiation (LTP) was induced by theta-burst stimulation (four trains of four pulses at 100 Hz; 5-Hz inter-burst interval) delivered at 0.1 Hz. LTP magnitude was calculated as the average (normalized to baseline) of the responses recorded 50–60 min after conditioning stimulation.

### Histological analysis

Fixed brains were collected as serial coronal sections, either on a freezing cryostat or embedded in paraffin and processed on a microtome, at 25-μm or 12-μm thickness, respectively (10 sections per stain per animal, from lambda 0 mm to lambda −4 mm). For the paraffin sections, a dehydration step was performed after staining. After formic acid-induced epitope retrieval, primary antibody 4G8 was incubated overnight at a 1:1000 dilution at room temperature (RT) (Biolegend, San Diego, CA). HRP-linked secondary goat anti-mouse antibody at a 1:1000 dilution (Invitrogen) was incubated for 2h at RT. The peroxidase reaction was visualized using a DAB kit (Vector) following the manufacturer’s instructions. Finally, sections were dehydrated through an ascending ethanol series, cleared in xylene, and coverslipped with DPX mounting medium (Innogenex, San Ramon, CA). For fibrillar Aβ quantification, sections were incubated in thioflavin S (ThS) solution (0.025% in 50% ethanol) for 10 min after thawing and coverslipped with DPX mounting medium (Innogenex, San Ramon, CA). All samples were analyzed on an inverted epifluorescence microscope (Olympus IX71), and quantification was performed using ImageJ software, measuring the area of fluorescent signal per total area (three images per animal for the brain cortex and two images for the hippocampus), after subtracting background signal.

### AβOs preparations

AβOs were prepared weekly from synthetic Aβ1–42 (California Peptide, Salt Lake City, UT, USA) and were routinely characterized by size-exclusion chromatography (SEC-HPLC) under non-denaturing conditions and, occasionally, by Western immunoblotting and transmission electron microscopy, as previously described^63^. Briefly, the peptide was dissolved in HFIP to 1 mM and stored as a dried film at -80°C after solvent evaporation. The film was resuspended in DMSO to a final concentration of 5 mM and thoroughly vortexed. The solution was then diluted in ice-cold PBS to 100 μM and left at 4°C overnight. The solution was centrifuged at 14,000 × g for 10 min at 4°C to remove insoluble aggregates (protofibrils and fibrils), and the supernatant containing Aβo was collected. Protein concentration was determined using the BCA assay (Thermo-Pierce). Intrahippocampal infusion of AβOs in mice was performed as described^64^.

### Stereotaxic injections

Sixty-day-old 5xFAD mice were anesthetized using isoflurane and fixed to a mouse stereotaxic frame (David Kopf Instruments). Bilateral injections of 2.5 μL of AAV-TT-XBP1s or PBS were performed at a single point in the hippocampal region (concentration stock 1x10^13^ vector genomes (Vg)/ml) using a 5-μL Hamilton syringe (Hamilton) at the following coordinates: AP: -1.8 mm, ML: 1.8 mm, and DV: -1.8 mm. The injection was delivered at a rate of 0.5 μL/min, and the needle was left in place for 5 min before retraction. During the stereotaxic procedure, the animals were injected subcutaneously with 5 mg/kg ketoprofen; they were then treated orally with ketoprofen for three consecutive days at the same dosage and monitored for 10 days following surgery using the Mouse Grimace Scale parameters ^65^.

### Real-time PCR analysis

Total tissue RNA was extracted using TRIzol™ Reagent (Invitrogen) according to the manufacturer’s instructions. cDNA was synthesized with random primers using a High-Capacity cDNA Reverse Transcription Kit (Applied Biosystems) and subsequently subjected to quantitative PCR analysis using HOT FIREPol® EvaGreen® qPCR Mix Plus (ROX) (Solis BioDyne) on a Stratagene Mx3000P instrument (Agilent Technologies). Actin mRNA expression was used to normalize all samples. The primer sequences used were as follows:

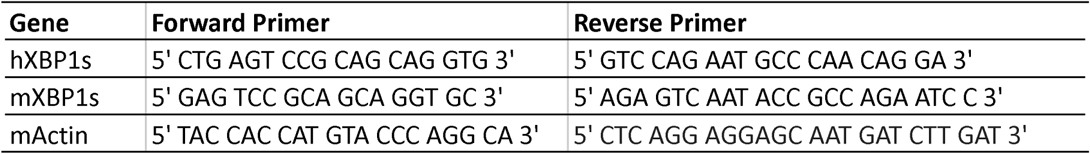

### Western blot analysis

Hippocampus and cortex samples were obtained from mice and lysed in PBS with a protease and phosphatase cocktail. Samples were then subjected to SDS-PAGE electrophoresis (12%), and proteins were transferred to activated PVDF membranes, blocked with 4% BSA for 1 h, and incubated overnight at 4°C with specific primary antibodies (XBP1s, BioLegend 658802, 1:250; Actin-HRP, Cell Signaling 5125S, 1:1000; and GFP, Abcam ab6556, 1:1000). Membranes were washed with 1× PBS, incubated with secondary antibodies, and protein bands were visualized using a Western ECL kit and a ChemiDoc™ XRS+ imaging system.

### Statistical analyses

Data are expressed as means ± SEM. Normality was assessed with the skewness/kurtosis test. Histological comparisons between two groups were analyzed with the Mann-Whitney U test. Behavioral and electrophysiological experiments with a factorial design (genotype × treatment, or AβO × treatment) were analyzed by two-way ANOVA with the interaction term as the test of treatment efficacy, followed by a pre-specified post-hoc comparison set with Bonferroni correction; learning curves were analyzed by repeated-measures two-way ANOVA with animal as the unit of analysis. Exact p values, effect sizes and 95% confidence intervals. Statistical analyses were performed using GraphPad Prism 5.0 software. Differences were considered statistically significant at p < 0.05.

## Acknowledgements

The authors thank María José Altamirano for her veterinary assistance. This work was funded by FONDECYT [1220573], ECOS- ANID [ECOS230024], ANID/FONDEF ID1ID22I10120, and US Army Medical Research Acquisition Activity (USAMRAA) AL2201415, HT9425-23-1-0990 (CH), ANID/NAM22I0057 (MD and CH), and ANID/FONDECYT 3190596 (MD).

## Conflict of interest

CH is co-inventor in different patent applications to use AAV2-XBP1s to treat ALS, memory loss and normal brain aging that were licensed to UCB, Belgium (PCT/CL2016/000056, WO2017059554A1, WO2016106458, U.S. Patent Application No.: 63/631,111).

## Supplementary figures

**Figure S1.**
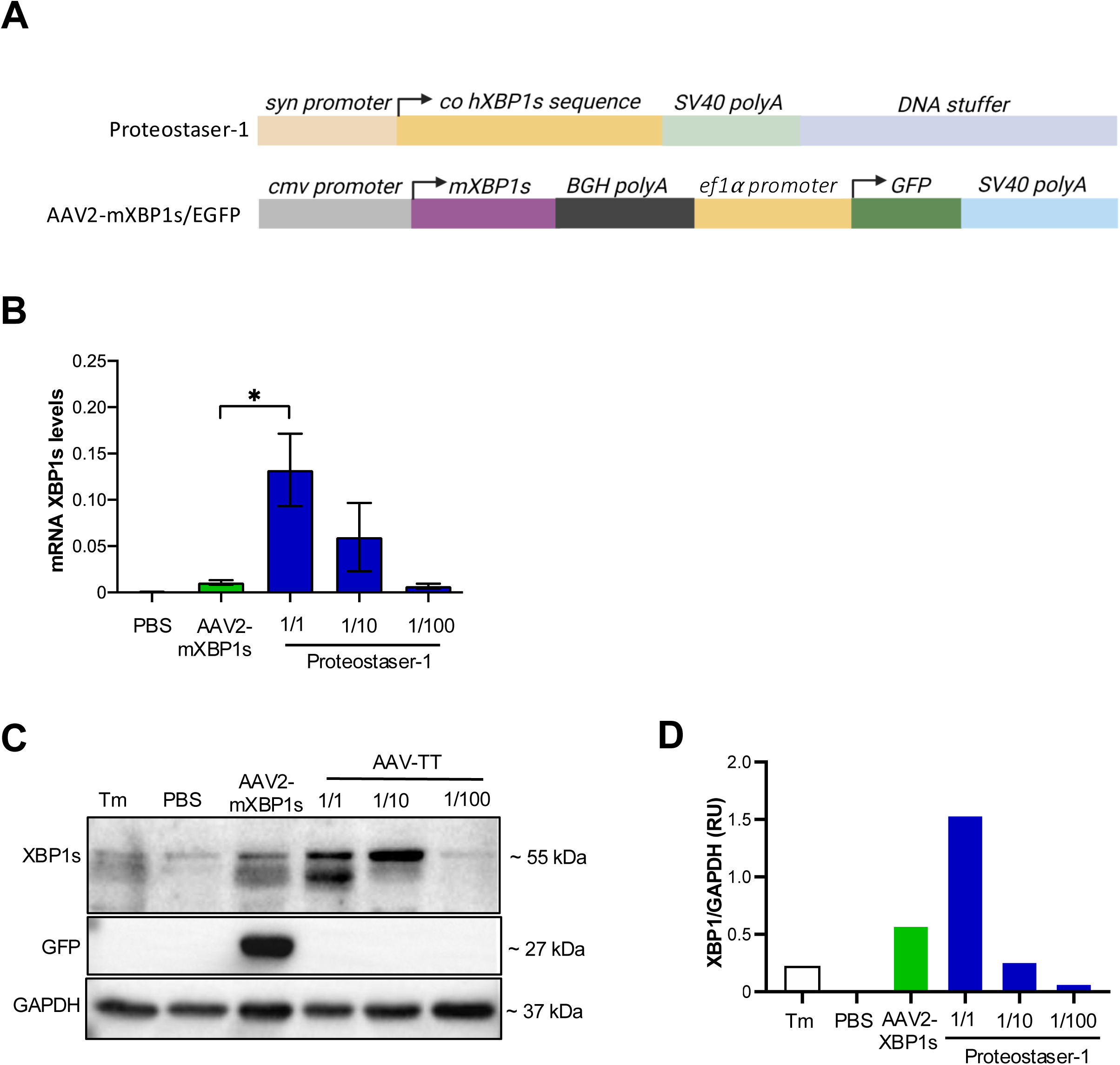
Generation of the Proteostaser-1 construct. (A) Scheme indicating the structure of the AAV2-mXBP1s/EGFP and AAV-TT-Syn-hXBP1s (Proteostaser-1) constructs. AAV2-mXBP1s/EGFP is an AAV serotype 2 construct that expresses mouse XBP1s cDNA under the control of the *CMV* promoter, in addition to containing an expression cassette for EGFP controlled by the *EIF2* promoter. Proteostaser-1 consists of an AAV-TT variant particle that expresses codon-optimized human XBP1s cDNA under the control of the mini synapsin (*syn*) promoter. (B) To compare relative expression levels, animals were injected with AAV2-mXBP1s/EGFP (2.5 μl x10^13^ Vg/ml per hemisphere) or indicated dilutions of Proteostaser-1 of the same concentration into the hippocampus using brain stereotaxis. One month later, hippocampus was isolated and *Xbp1s or XBP1s* mRNA levels were measured by real tome RT-PCR using primers that recognize human and mouse XBP1s cDNA. β-actin was used as a housekeeping gene to normalize gene expression. (C) Animals were injected with AAV2-mXBP1s/EGFP (n = 5), Proteostaser-1 at three concentrations (n = 5 per group), or PBS (n = 5) as indicated in B. One-way ANOVA followed by Tukey’s test was performed to confirm statistically significant differences (*: *p* < 0.001). (D) Left panel: XBP1s levels were determined in total protein extracts obtained from isolated hippocampus of animals described in B using Western blot analysis. In addition, as positive control, 5 mg/kg tunicamycin (Tm) was injected intraperitoneally to induce the UPR in the brain. GFP expression was also monitored (only expressed in AAV2-mXBP1s/EGFP), along with GAPDH as a loading control. Right panel: Relative XBP1s expression levels were quantified from Western blots shown in C.

**Figure S2.**
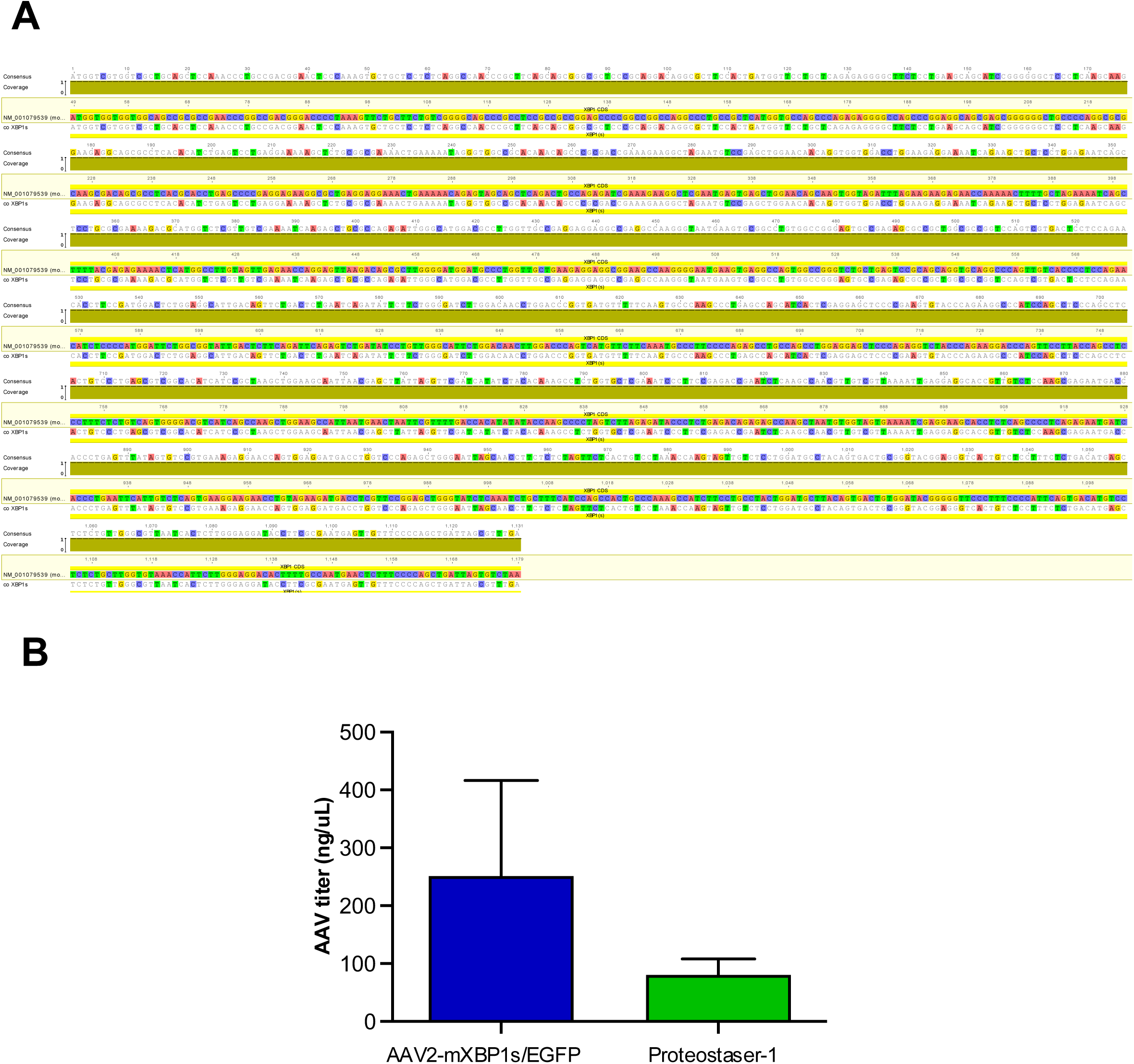
Generation of the codon-optimized human XBP1s AAV construct. (A) cDNA sequence of human XBP1s and the codon-optimized sequence used in the current study. (B) Comparative graph of single-strand DNA concentration determination for AAV2-mXBP1s/EGFP and AAV-TT-Syn-hXBP1s (Proteostaser-1). Sequence alignment is shown comparing the murine XBP1s reference sequence (NM_001079539) with the codon-optimized human XBP1s (co XBP1s) sequence used in Proteostaser-1. (B) Comparative graph of AAV titers determined by single-stranded DNA (ssDNA) concentration side by side for AAV2-XBP1s/EGFP and AAV-TT-Syn-hXBP1s

**Figure S3.**
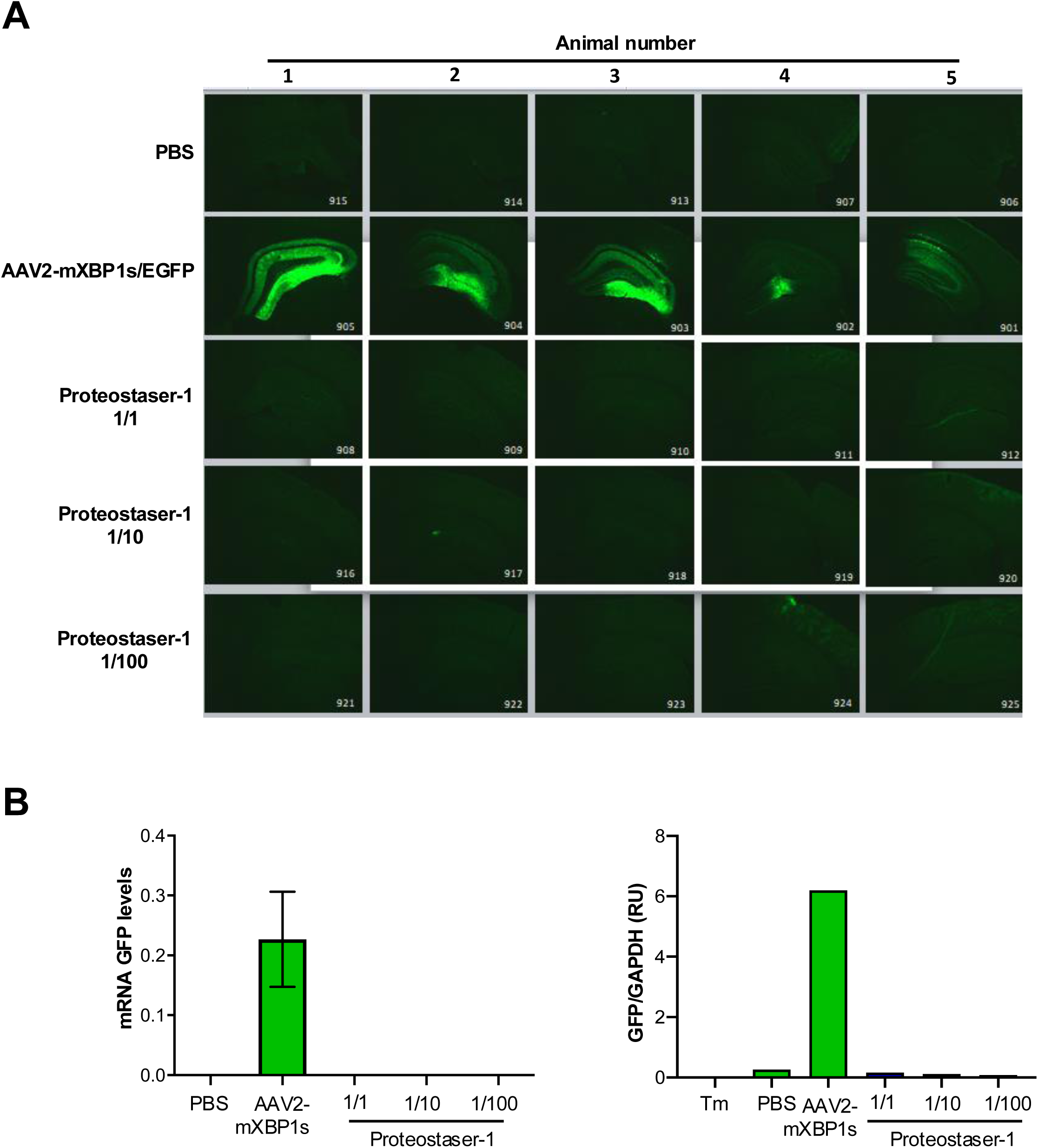
Transduction in the hippocampus of wild-type animals injected with AAV2-mXBP1s/EGFP or Proteostaser-1. (A) 2-month-old wild-type animals were injected with AAV2-mXBP1s/EGFP, Proteostaser-1 (1/1, 1/10, and 1/100), or PBS as control (n = 5 per group). After 1 month, animals were euthanized, and the presence of green fluorescent protein (GFP) in brain was assessed in histological sections. (B) Relative GFP expression in the hippocampus was measured by qPCR (n = 5 per group). Mean and standard error are shown. Statistical analysis was performed using one-way ANOVA followed by Tukey’s test to confirm statistically significant differences *: *p* < 0.05. (C) Western blot analysis to detect green fluorescent protein (GFP) in hippocampal protein extracts from 2-month-old wild-type animals injected with AAV2-mXBP1s/EGFP, Proteostaser-1 (1/1, 1/10, and 1/100), or PBS as control. As a positive control, 5 mg/kg of tunicamycin ™ was injected into wild-type animals.

**Figure S4.**
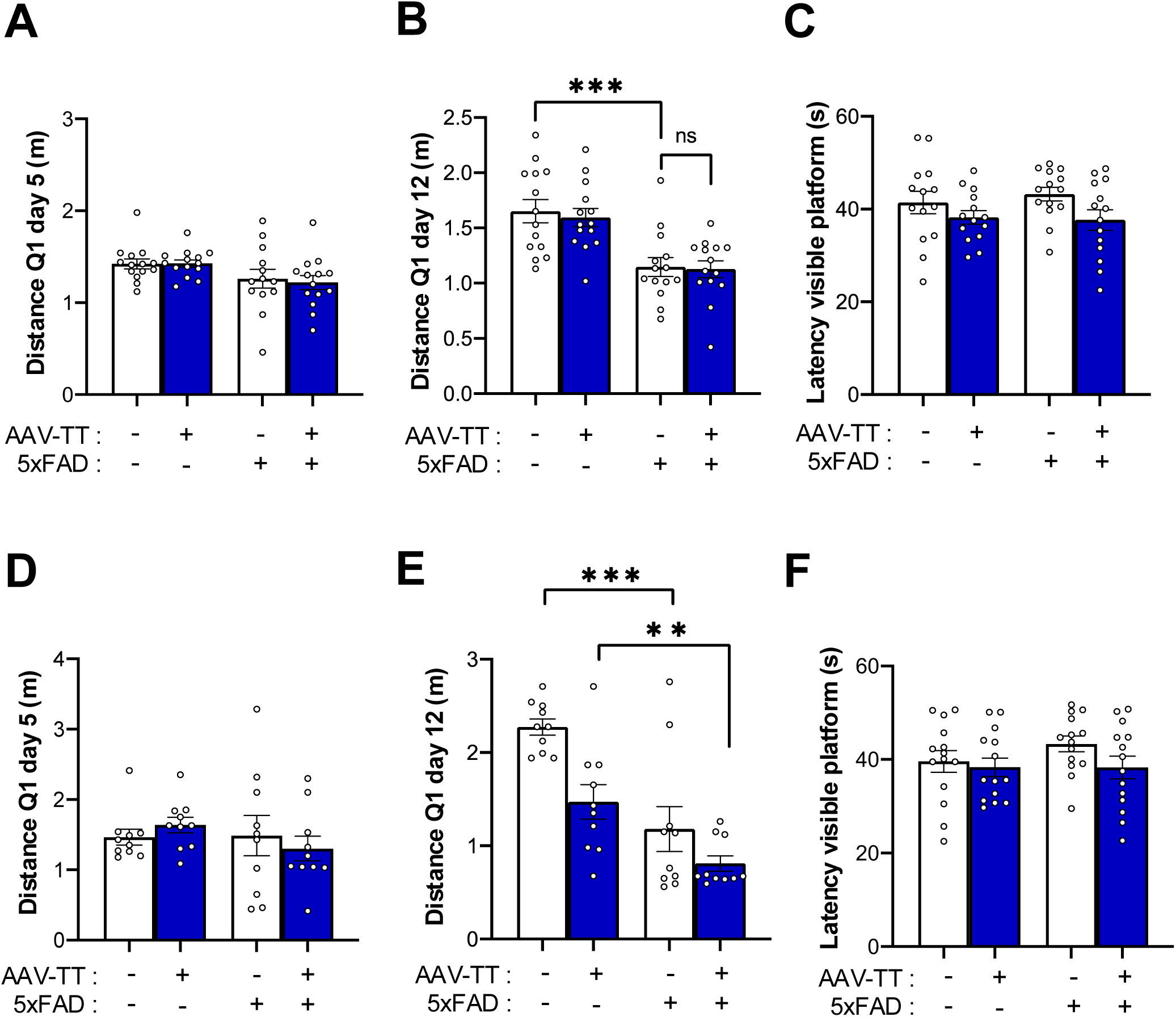
Assessment of behavioral performance in Proteostaser-1 treated 5xFAD mice. Two-month-old 5xFAD and non-transgenic littermate control mice received stereotaxic injections of Proteostaser-1 or PBS. Behavioral performance was assessed in the Morris water maze at 6 months (A–C) and 9 months of age (D–F). Probe trials were performed on days 5 (A, D) and 12 (B, E) after training to evaluate short- and long-term spatial memory, respectively, by measuring the distance traveled in the target quadrant (Q1) after removal of the platform. Following the day 12 probe trial, a visible platform test was performed to assess visual and motor function by measuring the latency to reach the visible platform (C, F). Mean and standard error are shown. Statistical analysis was performed using one-way ANOVA. *: *p* < 0.05; **: *p* < 0.01; ***: *p* < 0.001.

## Notes

### Competing Interest Statement

The authors have declared no competing interest.

